# A murine model of cardiovascular-kidney-metabolic syndrome demonstrates compromised limb function in the ischemic hind limb

**DOI:** 10.1101/2024.12.03.626707

**Authors:** Saran Lotfollahzadeh, Herreet Paul, Joshua Bonifacio, Ricardo Almiron, Isaac Hockestra, Kylla Prezkop, Trent Yamamoto, Maria Carmen Piqueras, Wenqing Yin, Marina Malikova, Jeffrey J. Siracuse, Mostafa Belghasem, Howard Cabral, Nazish Sayed, Vipul Chitalia

## Abstract

**Background:** Cardiovascular-Kidney-metabolic (CKM) syndrome is a public health problem involving > 90% of the US and results in premature CVD at a relatively preserved GFR. The molecular mediators of CKM are poorly understood, partly due to the lack of a reliable animal model. We set out to generate an animal model with renal and metabolic dysfunctions, using PAD as a CKM manifestation.

**Methods:** A group of C57BL/6 mice was randomized into four groups-a normal diet (ND), a 0.2% adenine diet (AD, a CKD model), a high-fat diet (HFD, a metabolic model), and a combination of HFD+AD (a potential CKM model). Mice underwent a hind limb ischemia surgery. An array of structural (capillary density, soleus muscle evaluation), and functional assays were performed.

**Results:** Compared to ND mice, AD mice showed loss of weight (40%) and GFR (90%) (P<0.001). The HFD+AD mice had 23-50% higher weight and GFR than the AD group (P = 0.003). The kidneys of HFD+AD showed tubular atrophy, tubulointerstitial fibrosis, immune infiltration, glomerulomegaly, consistent with glomerular hyperperfusion, hypercholesterolemia, impaired glucose tolerance, and adipophilin in the liver, an early marker of hepatic steatosis, and myocardial fibrosis. Compared to ND mice, the AD and HFD+AD mice showed similar reductions in the hind limb perfusion ratios, microcapillary density, Type II muscle fibers, and increased muscle fibrosis, immune infiltration, and lowest cross-sectional muscle area. Compared to AD, HFD+AD mice showed a lower flux ratio and grip strength, all at a GFR double that of the AD group.

**Conclusion:** A combination of HFD+AD in mice displays features of CKD (loss of GFR, renal fibrosis), dysfunctional obesity (dyslipidemia, impaired glucose tolerance, hepatic steatosis and glomerulomegaly), and cardiovascular disease (myocardial fibrosis and PAD) at a higher GFR, consistent with the features of CKM. This model can be explored to probe the mechanisms of CKM syndrome.

## Introduction

The interactions between chronic kidney disease (CKD), metabolic disorders (obesity and diabetes), and cardiovascular disease are well established. Recently, these morbidities have been considered holistically as Cardiovascular-Kidney-Metabolic (CKM) syndrome^1, 2^. CKM is defined as a health disorder due to connections among heart disease, CKD, diabetes, and obesity, leading to earlier presentation of CVD and poor health outcomes at a relatively preserved renal function^1,2^.

The CKM is recognized due to an increasing burden of obesity, diabetes, and CKD in the US^3^. Studies have shown that severe obesity with a BMI of 40 to 45 reduces survival by 8 to 10 years, diabetes by 13 to 40 years, and stage 4 and 5 CKD by more than 20 years^4^. These patients suffer from a combination of these comorbidities (multimorbidities). A nationwide sample showed 10-year mortality rates for diabetes alone was 7.7%, CKD was 11.5%, and a combination of diabetes and CKD was ∼ to double the sum of its parts (31.1%)^4^. There is a change in the trend of CVD in CKM in the population^3, 5^. The National Health and Nutrition Examination Survey from 1988 to 2018 identified ten cardiometabolic and renal phenotypes. The prevalence of the ‘high blood pressure’ and ‘high cholesterol’ phenotypes decreased over time, contrasted by a rise in the ‘obesity’ and low eGFR’ phenotypes.

Despite its profound public health implications, the mediators of CKM, especially for prevalent phenotypes of low GFR and dysfunctional obesity, remain poorly defined. A dearth of reliable animal models of CKM partially drives this limitation. Although CKD rodent models are well established and represent different types of kidney disorders^6^, they do not demonstrate obesity or occasionally have an opposite phenotype, such as weight loss (in an adenine-induced CKD model).

One of the manifestations of CKM is peripheral artery disease (PAD)^7^. PAD affects over 230 million adults worldwide and is linked to an increased risk of adverse clinical outcomes, including other CVDs, as well as adverse limb events like gangrene and amputation^8^. Our previous work has shown that CKD significantly affects post-ischemic angiogenesis in a rodent model^9^. However, only CKD was investigated as an individual risk factor in this work. Therefore, we set out to generate an animal model using PAD as a manifestation of CKM. In this model, we combined diet-induced obesity and CKD with a hypothesis that such a model will emulate CKM syndrome with metabolic abnormalities and CVD at an earlier stage of CKD (a hallmark of CKM syndrome in humans).

## Methods

All the experiments were performed with IACUC IPROTO202200000072 at Boston Medical Center and Boston University School of Medicine.

### HLI

A detailed procedure is described in the past^9^. Following anesthesia, a 5-mm longitudinal incision, perpendicular to the inguinal crease, was made, and the femoral artery was ligated with a non-absorbable 5.0 silk ligature just distal to the origin of the Profunda Femoris. A second ligature was placed 5 mm distal to the superficial epigastric artery. The interval segment of the femoral artery was excised and the profunda femoris and circumflex branches were ligated and excised.

### Animal models

A group of 8-to-12 weeks-old C57BL/6 male mice (Jackson’s lab, catalog 000664) was randomized into four groups. They were fed a casein diet for a week, followed by casein-based purified diets (see **Table 1** for the composition of the diet, Research Diets, NJ, USA) for six consecutive weeks to induce different disease models. CKD was induced using a 0.2% adenine diet (AD), a model characterized by profound tubulointerstitial fibrosis and retention of uremic solutes like the patients with ESKD^9, 10^. Obesity was induced using a high-fat diet (HFD). Mice on a casein-based diet (normal diet-ND) served as a control. A combination of HFD + AD was investigated as CKM syndrome. For the HLI model, diets were initiated for one week before HLI surgery and were continued for an additional four weeks.

**Table 1.** Diet composition.

| Components | HFD |  | HFD+AD |  | ND |  | AD |  |
| --- | --- | --- | --- | --- | --- | --- | --- | --- |
|  | gm% | kcal% | gm% | kcal% | gm% | kcal% | gm% | kcal% |
| Protein | 26.2 | 20.0 | 26.2 | 20.0 | 19.2 | 20.0 | 19.2 | 20.0 |
| Carbohydrate | 26.3 | 20.1 | 26.3 | 20.1 | 67.3 | 70.0 | 67.2 | 70.0 |
| Fat | 34.9 | 59.9 | 34.8 | 59.9 | 4.3 | 10.0 | 4.3 | 10.0 |
| Total |  | 100.0 |  | 100.0 |  | 100.0 |  | 100.0 |
| kcal/gm | 5.2 |  | 5.2 |  | 3.8 |  | 3.8 |  |
| Ingredient (kg) | gm | kcal | gm | kcal | gm | kcal | gm | kcal |
| Casein | 200.0 | 800.0 | 200.0 | 800.0 | 200.0 | 800.0 | 200.0 | 800.0 |
| L-Cystine | 3.0 | 12.0 | 3.0 | 12.0 | 3.0 | 12.0 | 3.0 | 12.0 |
| Corn Starch | 0.0 | 0.0 | 0.0 | 0.0 | 506.2 | 2024.8 | 506.2 | 2024.8 |
| Maltodextrin | 125.0 | 500.0 | 125.0 | 500.0 | 125.0 | 500.0 | 125.0 | 500.0 |
| Sucrose | 68.8 | 275.2 | 68.8 | 275.2 | 68.8 | 275.2 | 68.8 | 275.2 |
| Cellulose, BW | 50.0 | 0.0 | 50.0 | 0.0 | 50.0 | 0.0 | 50.0 | 0.0 |
| Soybean Oil | 25.0 | 225.0 | 25.0 | 225.0 | 25.0 | 225.0 | 25.0 | 225.0 |
| Lard | 245.0 | 2205.0 | 245.0 | 2205.0 | 20.0 | 180.0 | 20.0 | 180.0 |
| Mineral Mix S | 10.0 | 0.0 | 10.0 | 0.0 | 10.0 | 0.0 | 10.0 | 0.0 |
| DiCalcium Ph | 13.0 | 0.0 | 13.0 | 0.0 | 13.0 | 0.0 | 13.0 | 0.0 |
| Calcium Carb | 5.5 | 0.0 | 5.5 | 0.0 | 5.5 | 0.0 | 5.5 | 0.0 |
| Potassium Cit | 16.5 | 0.0 | 16.5 | 0.0 | 16.5 | 0.0 | 16.5 | 0.0 |
| Vitamin Mix V | 10.0 | 40.0 | 10.0 | 40.0 | 10.0 | 40.0 | 10.0 | 40.0 |
| Choline Bitart | 2.0 | 0.0 | 2.0 | 0.0 | 2.0 | 0.0 | 2.0 | 0.0 |
| Adenine | 0.0 | 0.0 | 1.6 | 0.0 | 0.0 | 0.0 | 2.1 | 0.0 |
| FD&C Yellow | 0.0 | 0.0 | 0.1 | 0.0 | 0.0 | 0.0 | 0.0 | 0.0 |
| FD&C Red Dy | 0.0 | 0.0 | 0.0 | 0.0 | 0.0 | 0.0 | 0.1 | 0.0 |
| FD&C Blue D | 0.1 | 0.0 | 0.0 | 0.0 | 0.0 | 0.0 | 0.0 | 0.0 |
| Total | 773.9 | 4057.2 | 775.4 | 4057.2 | 1055.1 | 4057.0 | 1057.2 | 4057.0 |
| % Adenine |  |  | 0.2 |  |  |  | 0.2 |  |

### IHC and IF and quantification of features

For immunohistochemistry (IHC) and immunofluorescence (IF), tissue samples were processed = as previously described^10^. Antibodies are listed in **Table 2**. For image quantification, the slides were scanned using an automated motorized stage system with Nikon NIS Elements software at the Imaging Core. Images were analyzed in ImageJ software, where the signals were converted to grayscale, and pixel number and intensity were measured as integrated density using the Fiji plugin as done previously. The cross-sectional area (CSA) was designated as the region of interest and areas was calculated using ImageJ. The integrated density of each image was normalized to the respective area as done in the past^9–11^.

**Table 2.**
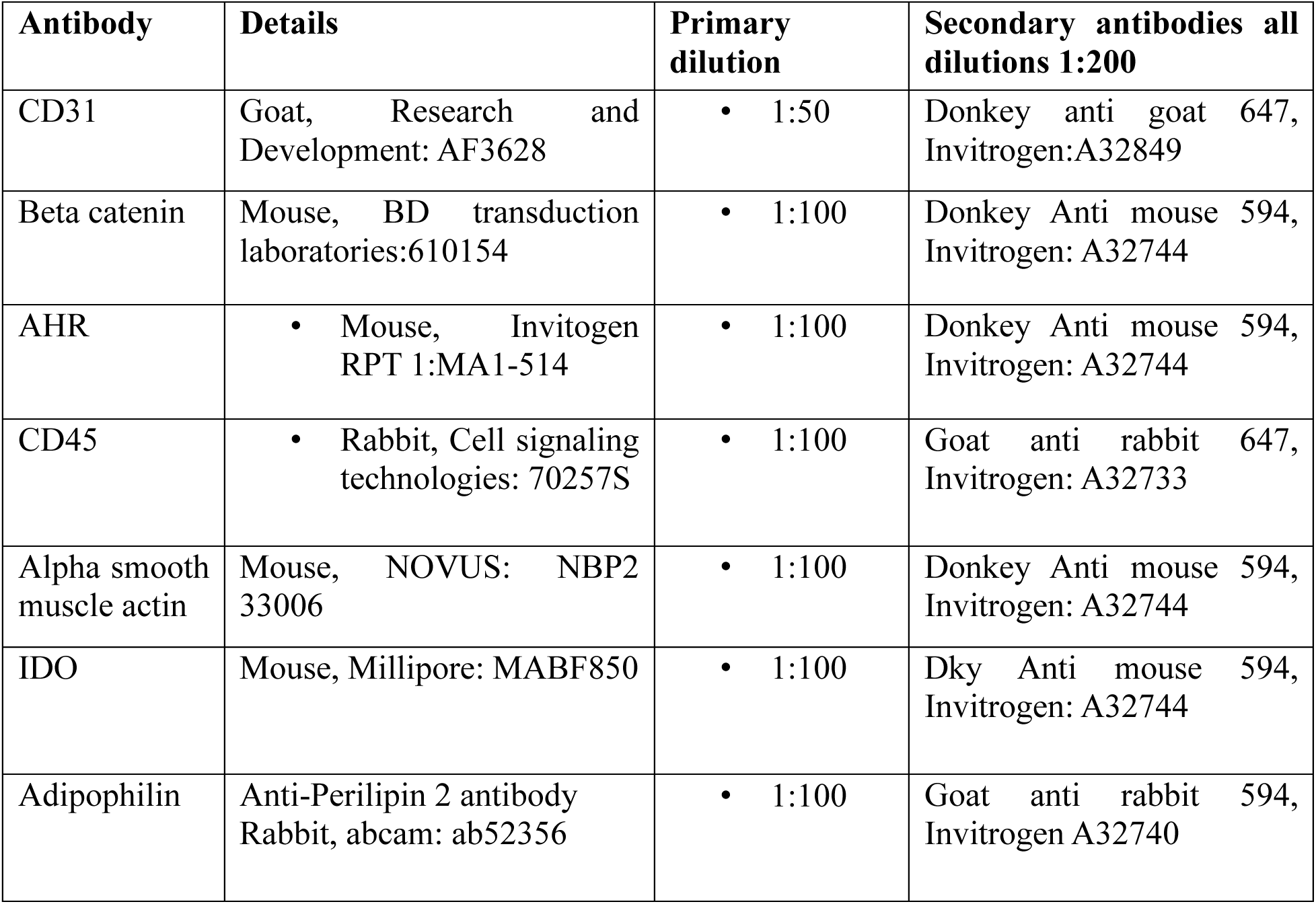
Antibodies used in this work.

| **Antibody** | **Details** | **Primary dilution** | **Secondary antibodies all dilutions 1:200** |
| --- | --- | --- | --- |
| CD31 | Goat, Research and Development: AF3628 | • 1:50 | Donkey anti goat 647, Invitrogen:A32849 |
| Beta catenin | Mouse, BD transduction laboratories:610154 | • 1:100 | Donkey Anti mouse 594, Invitrogen: A32744 |
| AHR | • Mouse, Invitrogen RPT 1:MA1-514 | • 1:100 | Donkey Anti mouse 594, Invitrogen: A32744 |
| CD45 | • Rabbit, Cell signaling technologies: 70257S | • 1:100 | Goat anti rabbit 647, Invitrogen: A32733 |
| Alpha smooth muscle actin | Mouse, NOVUS: NBP2 33006 | • 1:100 | Donkey Anti mouse 594, Invitrogen: A32744 |
| IDO | Mouse, Millipore: MABF850 | • 1:100 | Dky Anti mouse 594, Invitrogen: A32744 |
| Adipophilin | Anti-Perilipin 2 antibody Rabbit, abcam: ab52356 | • 1:100 | Goat anti rabbit 594, Invitrogen A32740 |

### Exercise test

Mice were trained to run on an Exer 3/6-RB Treadmill (Columbus Instruments, Columbus, OH). Before HLI, four training sessions were given over two weeks, with only one training session per 24-hour period. The treadmill was placed at a 10-degree incline, shocks were enabled and administered at 1Hz per second, and treadmill lanes were cleaned with 70% ethanol between mice. Training sessions consisted of mice walking on the treadmill at 10 meters/min for 10 minutes and then 5 meters/min for 5 minutes. After the 4 training sessions, exhaustion testing was performed once before HLI surgery and then once a week after surgery. Exhaustion testing consisted of mice warming up on the treadmill starting at 5 meters/min and then increasing speed by 1 meters/min every minute until 10 meter/min was reached; at this point, distance and time recording began. After 5 minutes, the speed was increased to 15 meters/min and then speed was increased by 3meter/min every 5 minutes until a top speed of 30 meters/min was reached. The test concluded when the mice reached exhaustion, defined as a mouse spending 10 continuous seconds on the shock mat. At this point shocks were disabled, and distance and time to exhaustion were recorded.

### Grip test

Grip strength was measured using a grip strength meter (Imada DS2-50N digital force gauge) set to peak force mode. Mice were held by the tail and placed such that only the hindlimbs grasped the bar. The animals were allowed to grab a mesh wire with their forepaws. The mice were gently pulled backwards away from the grip strength meter, and the peak force was recorded in gram-force (gf). Three measurements were taken during each session, with the mice being allowed to rest for at least 1 minute between measurements. The greatest of the three measurements was used as the result.

### Flux ratio and Doppler Laser

To assess perfusion in the hind limbs of the mice, a moor LDI Laser Doppler perfusion imaging instrument with software version 5.3 (Moor Instruments, Wilmington, DE) was used as described previously^9^.

### Glomerular filtration rate (GFR) measurement using FITC-Sinistrin preparation

We used a 10mg/100g dose of FITC-Sinistrin (Medibeacon FITC-Sinistrin) for GFR measurement via retro-orbital bolus per the manufacturer’s instructions^12^.

### Glucose intolerance test

An oral glucose load was injected per the protocol described previously^13–15^.

### Statistical analysis

Statistical analysis was conducted using SAS version 9.3 (SAS Institute). Data are presented as mean ± SEM, or as median with range and 25th and 75th percentiles in violin plots. Group comparisons were assessed using an ANOVA or two-tailed Student’s t-test. The linear mixed models were performed for analysis focused on the time-by-group interaction, comparing and testing for differences in the patterns of mean across groups. Finally, the post-hoc analysis was performed for the p-values for pairwise comparisons between groups at the final time point adjusted for multiple hypothesis testing at that time point. Statistical significance was defined as P ≤ 0.05.

## Results

### Characterization of kidneys in different groups of mice

The weights of mice in all four groups were recorded weekly (**Figure 1A**). A mixed linear model showed a significant interaction between time and group (P < 0.0001). The pre-randomization weights were similar in all groups. The mice on ND showed a minimal increase in weight over the length of the experiment. However, mice on HFD showed a persistent increase in body weight, which was higher by ∼ 20% compared to the ND group at the end of the experiment (ND mice = 26.23 +/- 1.7 grams, HFD 30.493 + 2.374 grams, P = 0.03). Mice on AD showed a consistent reduction in weight from ∼23 grams to 15.4 +/- 1.2 grams over the 6-weeks period (P<0.0001 compared to ND). Mice on AD + HFD had an increase in body weight in the first two weeks, followed by a reduction at the end of the experiment, with the final body weight lower than that of AD (HFD+AD = 22.46 +/-2.49 grams, P < 0.001) and HFD (P = 0.003).

**Figure 1.**
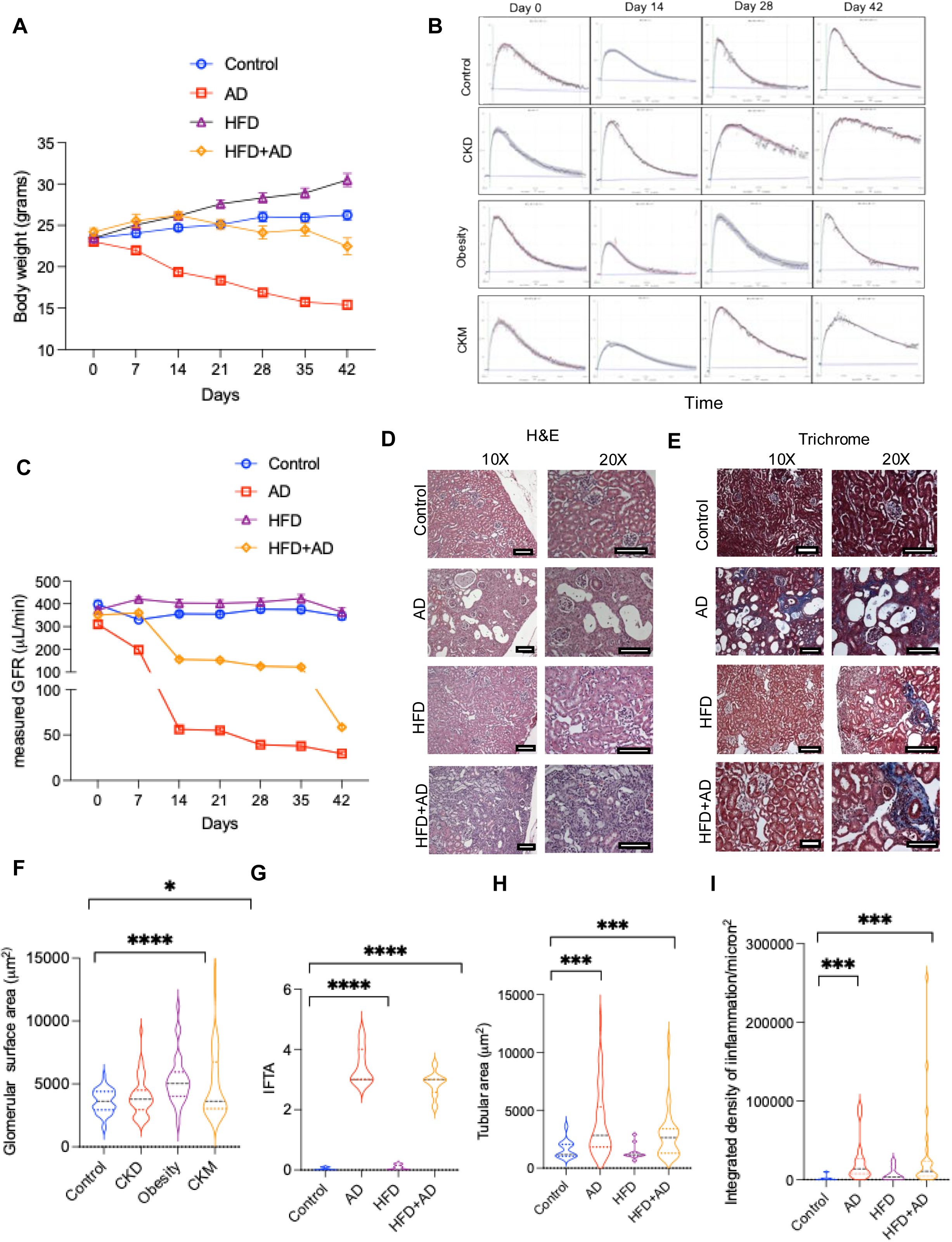
Characterization of kidneys of mice from four different groups. **(A)** The average weights of mice in all four groups are shown. Error bars-SEM. The mixed linear model was followed by a post-hoc analysis for different group comparisons, which were adjusted for multiple comparisons. **(B)** Representative graphs of glomerular Filtration Rate in mice measured using FITC-SINISTRIN dye per group are shown. **(C)** The average GFR of mice measured using FITC-Sinistrin dye is shown. The mixed linear model showed time and group interaction with P <0.0001, group P <0.0001, and time and group interaction P <0.0001. Tukey-Kramer adjustment was performed was multiple comparisons. **(D)** Kidneys were stained with hematoxylin and eosin from all four groups. Scale bar = 100 microns. **(E)** Representative images of kidneys stained with modified Masson Trichome are shown. Scale bar = 100 microns. **(F)** The glomerular surface area measurements on three high power field (hpf) images from each kidney of mice were performed ANOVA, P = 0.0002. Post-Hoc Student’s t-test was performed to compare the groups. ND and AD P = 0.1761. ND and HFD ****p < 0.0001, ND and HFD+AD P < 0.0001. **(G)** IFTA scores measured from three hpf images across four experimental groups of mice are shown. ANOVA analysis showed P < 0.0001. No significant difference was observed between ND and HFD groups. **(H)** Tubular area measurements across experimental groups. ANOVA P < 0.0001. Post-hoc pairwise comparisons using Student’s t test was performed. **(I)** The normalized integrated density of inflammation (per μm^2^) was obtained from three hpf per kidney per mouse in each group. ANOVA P = 0.2161. Post-hoc pairwise comparisons using Student’s t test were performed.

Given the change in the lean body weights in different groups, we measured GFR using the FITC-Sinistrin dye weekly (**Figure 1B**). Faster dye clearance suggests a higher GFR level, while delayed dye clearance points to a reduced GFR. Mice on the ND had GFR unchanged over 42 days, while mice on HFD showed persistently higher GFR (pre-harvest GFR ND = 345.51+/- 52.53 μL/min, HFD 363.75 +/-57.51, P = 0.03, **Figure 1C**). As expected, mice on AD showed a drastic reduction in GFR in the first 14 days, and it plateaued to be around 10% GFR at the beginning of the study (29.41+/-4.4 μL/min). Mice with HFD+AD followed a similar trend to AD mice. However, at each time point, the GFR values were greater in HFD+AD mice compared to mice on AD (pre-harvest 58.53 +/- 5.10 μL/min, P <0.0001, **Figure 1C**). At the end of the experiment, blood urea nitrogen in the ND group was 23.10 +/- 4.45 mg/dL; the AD group was 134.2 +/-7.24 mg/dL, HFD = 28.80 +/- 2.77, and HFD +AD was 96.60+/- 0.89 mg/dL. Creatinine was ND – 0.10 +/- 0, AD = 1.26 +/- 0.23, HFD = 0.30 +/- 0.27 and HFD +AD 0.72 +/- 0.08 mg/dL.

The kidneys of mice on AD showed global glomerulosclerosis, tubule-interstitial fibrosis, loss of brush border, dilated tubules, tubular atrophy, and immune cell infiltration. Mice on HFD showed glomerulomegaly and minimal immune infiltration. The HFD+AD mice had features of CKD along with glomerulomegaly (**Figure 1D-1E**). The kidneys of mice on AD and HFD+AD showed interstitial fibrosis (blue color on the trichrome stain) more compared to other groups. The morphological features of kidneys were quantitated by a nephropathologist blinded to the group of mice (M.B.). Compared to the ND mice, the glomerular surface area was higher in HFD (P<0.0001) and HFD + AD (P = 0.031) (**Figure 1F**). The interstitial fibrosis and tubular atrophy (IFTA) were quantitated by the intensity of intensity and distribution of fibrosis along with tubular features, as done previously^16^. A scale of 1 to 5 was used, where a score of 5 indicated IFTA affecting 100% of the renal parenchyma. Compared to ND, mice on AD and HFD+AD showed significantly higher IFTA (P<0.0001) (**Figure 1G**). The kidneys showed tubular dilatation due to tubular cell atrophy, which was higher in AD (P= 0.003) and HFD +AD (P = 0.005) groups compared to ND (**Figure 1H**). The immune infiltration was significantly higher in AD (P = 0.005) and HFD+AD (P= 0.0019) groups compared to ND (**Figure 1I**). The HFD+AD mice demonstrate changes consistent with a combination of glomerulomegaly, a pathognomic feature of obesity, and renal damage (tubular atrophy, interstitial fibrosis, reduced GFR).

### Characterization of metabolic derangements and cardiac alterations in different groups of mice

CKM is characterized by dyslipidemia, glucose intolerance, and evidence of hepatic steatosis. At the end of the experiment, compared to the ND group, serum cholesterol showed a trend towards higher levels in AD (P= NS), while HFD and HFD+AD showed a 20-40% increase in serum cholesterol levels. The HFD+AD showed 20-30% higher cholesterol than the HFD group alone. In contrast, triglyceride levels remained similar between groups (**Figures 2A** and **2B**). The ND and AD groups showed a similar glucose tolerance test, while HFD and HFD+AD showed an impaired glucose tolerance (**Figure 2C**). Adipophilin binds to cytoplasmic lipid droplets and is an early marker of hepatic steatosis^17^. Compared to mice on ND, livers of mice on HFD and HFD+AD showed ∼ doubling of adipophilin expression (**Figure 2D** and **2E**). Myocardium was stained for fibrosis and collagen deposition, the pathological features associated with heart pathologies, including heart failure^18^ (**Figures 2F-2H**). The hearts of AD and HFD+AD mice showed a significant increase in fibrosis (Trichrome stain blue color) and collagen deposition (on Sirius red stain) (**Figure 2F**) compared to the ND group. Taken together, mice on HFD+AD develop hypercholesterolemia, impaired glucose tolerance, and liver steatosis, demonstrating the metabolic derangements consistent with CKM syndrome, while the AD-induced CKD model lacked some of these features. Also, CKM mice showed early evidence of myocardial fibrosis.

**Figure 2.**
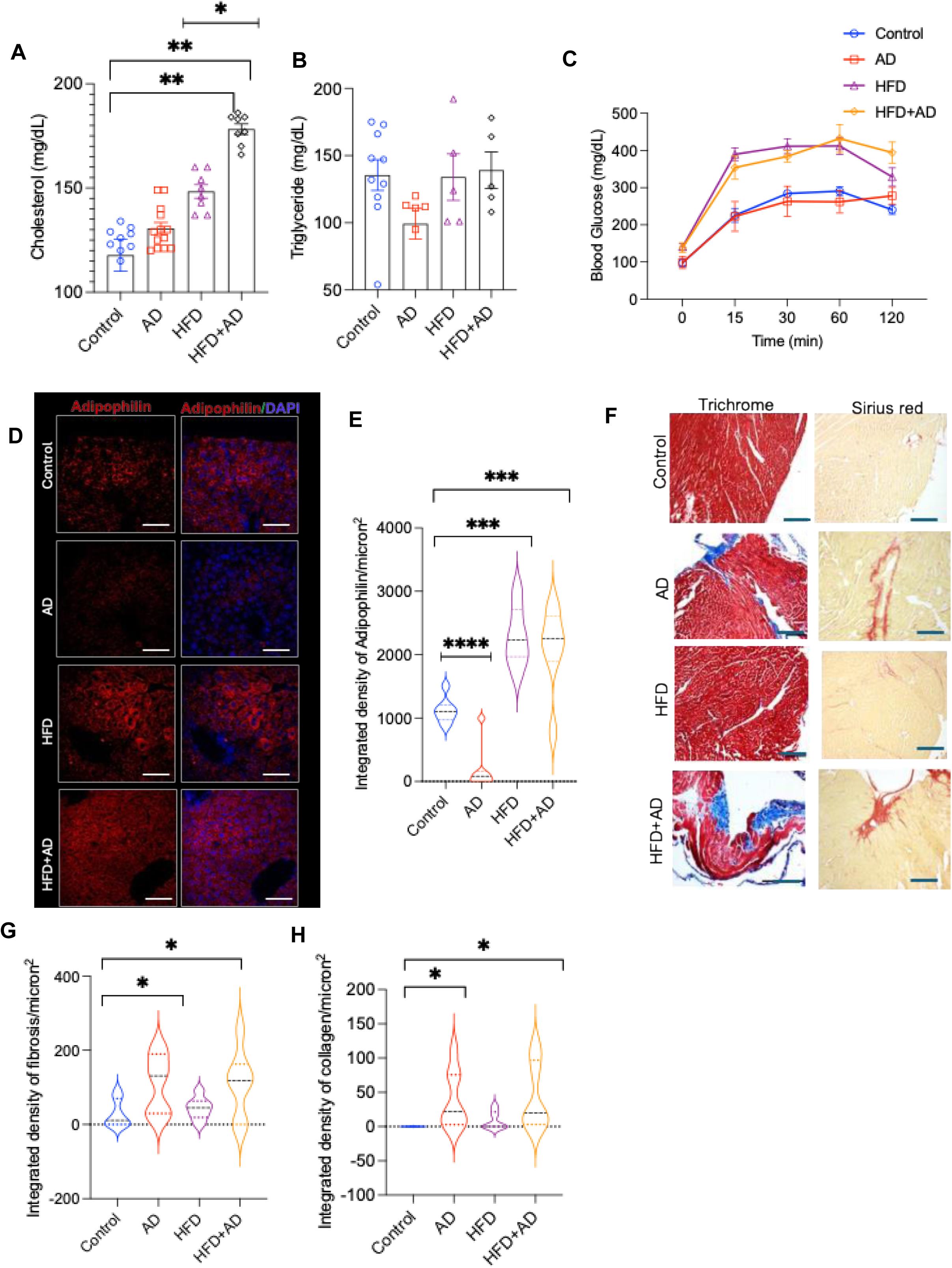
Characterization of metabolic parameters and heart morphology in four groups. **(A)** Average cholesterol levels are shown. Error bars-SEM. One-way ANOVA P <0.001. ND vs. HFD P = 0.003, ND vs. HFD+AD P = 0.002, and HFD vs. HFD +AD P = 0.0436. **(B)** Average triglyceride levels are shown. Error bars – SEM **(C)** Glucose tolerance test with average and SEM of glucose at indicated time points after glucose load are shown. ANOVA P < 0.001. At 15 minutes, blood glucose levels were significantly higher in HFD and HFD +AD groups compared to ND P < 0.0001. At 30 and 60 minutes, ND and both HF and AD groups P < 0.001 for both time points. At 120 minutes, ND and HFD (P = 0.039). No significant differences were observed between other group comparisons across all time points. **(D)** Representative liver sections stained for Adipophilin from four groups of mice are shown. Scale Bar = 100 microns. **(E)** Adipophilin expression was quantified in liver biopsies of all four groups as integrated density normalized to area. Average and SEM are shown. ANOVA < 0.0001. **** P < 0.0001, ***P = 0.0028. **(F)** Representative images of myocardium stained with modified Mason Trichrome (left column) and Sirius Red (right column) are shown. Scale bar = 100 microns. **(G)** The averages of myocardial fibrosis quantified from three hpfs of Trichrome-stained myocardium per mouse are shown as integrated density normalized to the surface area. ANOVA for all groups P-value of 0.0435. Compared to ND, AD group P = 0.0302, and HFD+AD P = 0.0451. **(H)** The averages of myocardial collagen quantified from three hpfs of Sirius Red-stained myocardium per mouse are shown as integrated density normalized to the surface area. ANOVA P-value of 0.0256. Compared to ND, the AD group P = 0.0193, and the HFD+AD group (P = 0.0219).

### PAD manifestation in different groups of mice

In the HLI model, perfusion recovery following femoral artery ligation serves as a biological readout of post-ischemic angiogenesis. It is presented as a ratio of perfusion in the ligated limb to the unligated limb indicating post-ischemic angiogenesis. For this experiment, mice were initiated on different diets for one week before HLI, and then the diets were continued through the study period. The pre-procedural perfusion ratios were ∼ 1 in both limbs and reduced to 10% of the pre-procedure values in all groups (**Figure 3A-3B**). In the ND group, the perfusion increased by 50% and 65% of the pre-procedural value at the end of the experiment (**Figure 3B**). The perfusion ratio was higher at day 14 after HLI in HFD and then plateaued. Compared to ND, the perfusion ratios of mice in HFD+ AD were ∼15% higher (P =NS) and then reduced at day 28 (P = 0.031). At the end of 28 days, compared to ND, both AD and HFD+AD showed ∼ 50% reduction in the perfusion ratio (P< 0.0001). There was no significant difference in the perfusion ratio between the AD and the HFD+AD group.

**Figure 3.**
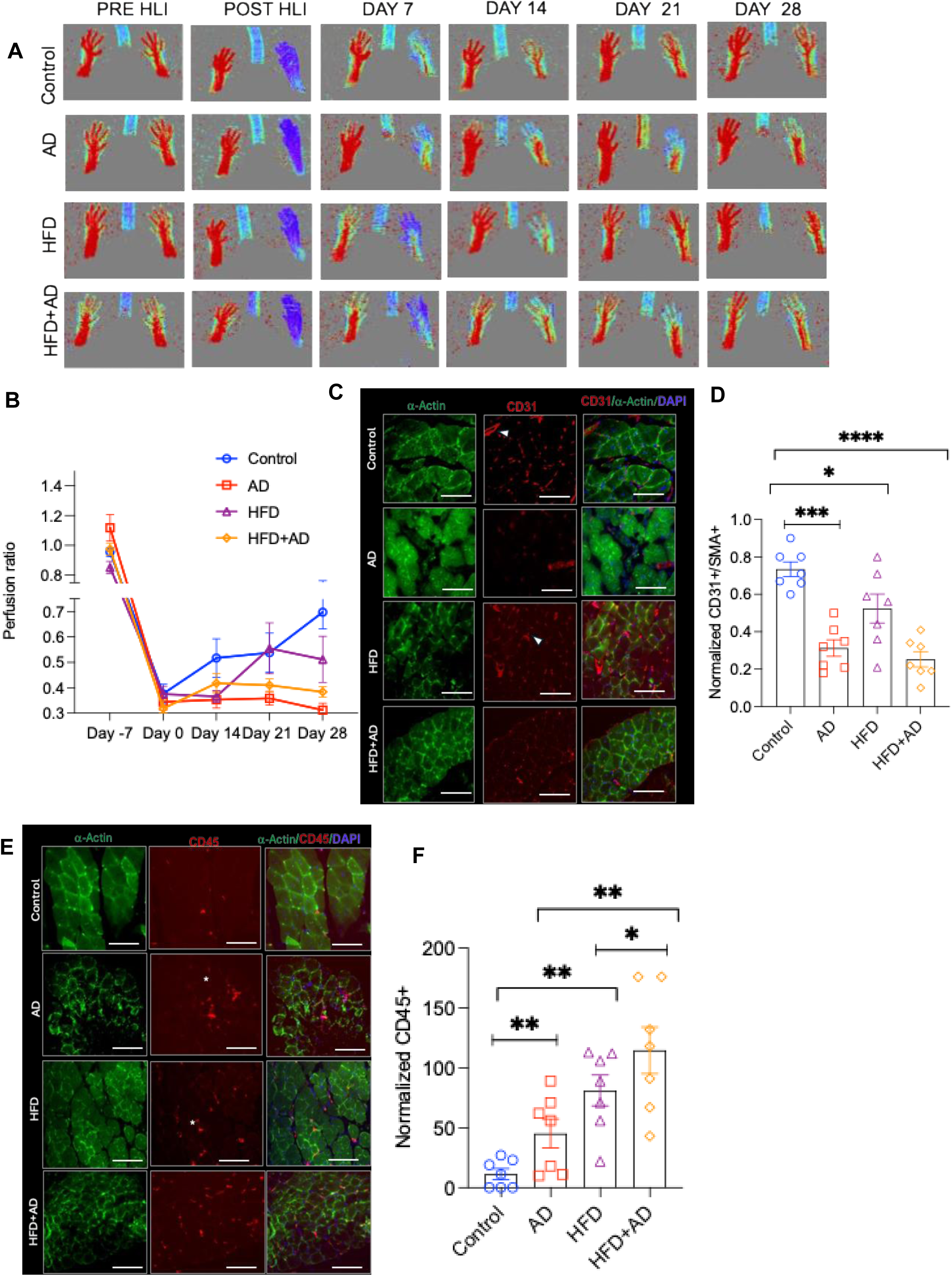
CKM mice demonstrate reduced microcapillary density and VEGF expression in the ischemic hind limbs. **A)** Representative laser Doppler images of mouse hind limb across four groups, among 6 times points of Pre-HLI, immediate post-HLI, Day 7, 14, 21, and 28 after HLI (N= 7 per group). **B)** The average and SEM of the perfusion ratio from all experimental groups are shown. A linear effects model showed a significant interaction between perfusion ratio and time P < 0.0001. Post hoc analysis with Tukey-Kramer adjustment indicated that at day 28, the ND vs AD P < 0.0001, ND vs HFD+AD group P = 0.0315. **C)** Three random hpf images from soleus muscles of the ischemic limb of each mouse (N =7 mice/group) stained with CD31 and α actin are shown. White arrowheads are directed at CD31+ capillaries. Scale bars = 100 microns. **D)** The average CD31 expression normalized to α actin was quantified as integrated density. One-way ANOVA *P* < 0.0001. Post hoc analysis was performed to compare groups. **E)** Three random hpf images from soleus muscles of the ischemic limb of each mouse (N =7 mice/group) stained with CD45 and α actin are shown. White asterisks correspond to CD45 expression. Scale bars = 100 microns. **F)** The average CD45 expression normalized to α actin was quantified as integrated density. One-way ANOVA *P* < 0.0001. Post hoc analysis was performed to compare groups. Compared to HFD, CD45 was greater in the HFD+AD group P = 0.043. Compared to the AD group, HFD+AD P = 0.004.

The perfusion to the lower extremity is based on the microcapillary development in skeletal muscles of the ischemic hind limb. The capillary density was examined in the soleus muscle of ischemic and non-ischemic limbs in all four groups using CD31, a marker of ECs, and normalized to α-actin^9^. As previously described, the expression of both α-actin and CD31 were quantitated as the integrated density^9, 10^. (**Figure 3C-3D**). Compared to ND, the normalized CD31 expression was reduced to 50% in AD (P<0.001), 15-23% in HFD (P= 0.0318), and 45% in the HFD+AD groups (P <0.0001). To probe the muscle immune microenvironment, the soleus was stained using a CD45 antibody, a marker of leucocytes. The ND mice showed minimal expression of CD45, while all the other groups of mice had a 30-80% higher expression of CD45 (AD P = 0.0220 and HFD P = 0.032) with the highest in HFD+AD mice (P = 0.002, **Figure 3E-3F**). Also, the HFD+AD group showed a higher CD45 expression than the AD and HFD groups. These results suggested that mice with HFD+AD showed compromised angiogenesis similar to AD mice at a GFR value ∼ double than that of the AD group (**Figure 1C**) and the highest immune cell infiltration compared to other groups.

### Characterization of myopathy in different groups of mice

Skeletal muscle damage in these mice can be an effect of compromised angiogenesis and a direct effect of CKD and metabolic factors. The cross-sectional areas (CSA) soleus muscles of non-ischemic and ischemic limbs were compared in four groups. On the non-ischemic limb, all groups showed a 10-15% reduction in CSA compared to the ND mice (**Figures 4A** and **4B**). The mean CSA of soleus of ND mice was 1200 µm^2^, and AD, HFD, and HFD+AD were 900, 1100, and 800 µm2, respectively. This phenotype was exacerbated by ischemia (**Figures 4C** and **4D**). Compared to the non-ischemic group, the ischemic limb soleus of all groups showed lower CSA (**Figures 4B** and **4D**). Compared to ischemic soleus from ND mice, AD had ∼ 50% reduction in CSA (P = 0.003), which was also seen in mice with HFD (P= 0.023). Mice with HFD+AD had the lowest CSA (500 µm^2^ P <0.0001), suggesting severe muscle atrophy in this group.

**Figure 4.**
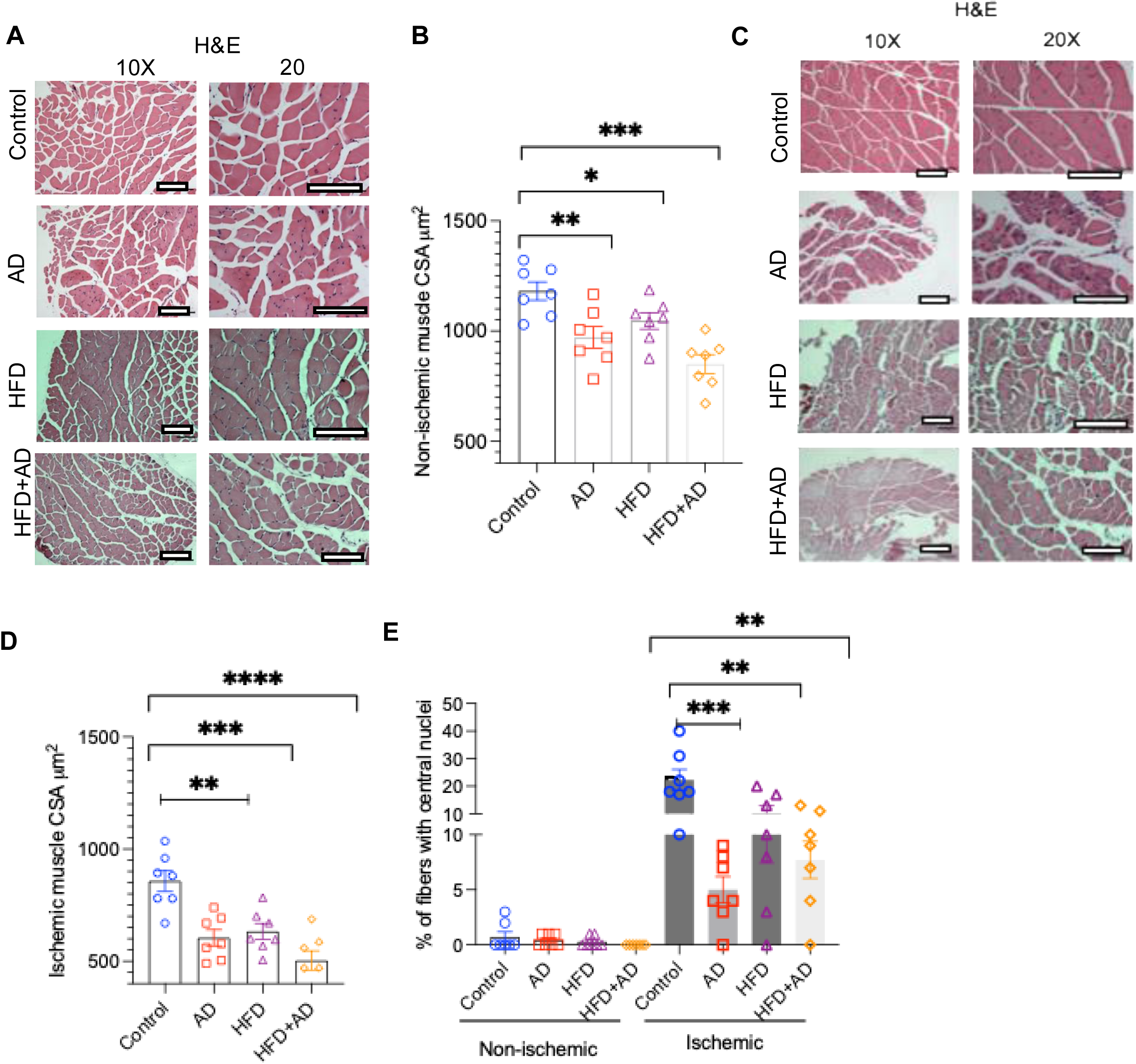
Characterization of myocytes in four groups of mice. **A)** Representative images of soleus muscle on the sham-operated limbs (non-ischemic) stained with H & E and obtained at two different magnifications are shown. Scale bar= 100 microns **B)** The average cross-sectional area (CSA) of non-ischemic soleus muscle obtained from five random high-power field images of each mouse are shown. Error bars - SEM. One-way ANOVA P = 0.0001. Compared to ND, the AD group P = 0.0067, the HFD group P = 0.0334, and the HFD+AD group P = 0.0001. **C)** Representative images of soleus muscle on the HLI-operated limbs (ischemic) stained as above. Scale bar= 100 microns **D)** The average CSA of the ischemic soleus muscle obtained from five random high-power field images of each mouse are shown. Error bars - SEM. One-way ANOVA P = 0.0001. **E)** Percentage of muscle fibers with central nuclei across experimental groups in the ischemic and non-ischemic muscle tissue. Three random hpf images per mouse were examined in a manner blinded to the group. Two-way ANOVA showed P < 0.0001. In non-ischemic muscles, no significant differences were observed. In the ischemic muscle, compared to ND, the AD group P = 0.0032, HFD P = 0.0043, and HFD+ AD P = 0.0395.

The ischemic insult is associated with the proliferation of skeletal muscles characterized by the central nucleus. As expected, the soleus on non-ischemic limbs showed an almost negligible number of proliferating fibers (**Figure 4E**). In the ischemic group, the percentage of muscle fibers with central nuclei in ND mice was ∼ 20% of total fibers, which was reduced by more than 80% in AD and by 50% in HFD and HFD+AD diets (Figure 4A), suggesting compromised regeneration of skeletal muscles in these groups.

Muscle atrophy is associated with fibrotic tissue. We further examined muscle fibrosis in both non-ischemic and ischemic limbs using a modified Masson Trichrome stain (**Figures 5A** and **5B**). In the non-ischemic group, the soleus of AD and HFD+AD showed an increase in fibrosis compared to ND controls (**Figure 5C**). The fibrosis was exacerbated in the ischemic group. In the ND group, ischemic soleus showed a 30% increase in fibrosis compared to the non-ischemic group. In the ischemic limb, compared to ND, mice in AD and HFD+AD groups showed a 60% to a 4-fold increase in fibrosis (**Figure 5C**). Soleus predominantly contains glycolytic type II fibers, which were identified using MYH7^19^. In the ischemic limb, compared to the ND group, type II fibers were reduced by 50-80% in AD and HFD+AD groups, respectively (**Figures 5D** and **5E**). The HFD+AD group showed 30% lower MYH7 compared to the AD group. These results demonstrate the compromised regenerative ability of the ischemic muscles and muscle fibrosis in all groups of mice and the highest muscle atrophy and immune infiltration in the HFD+AD group.

**Figure 5.**
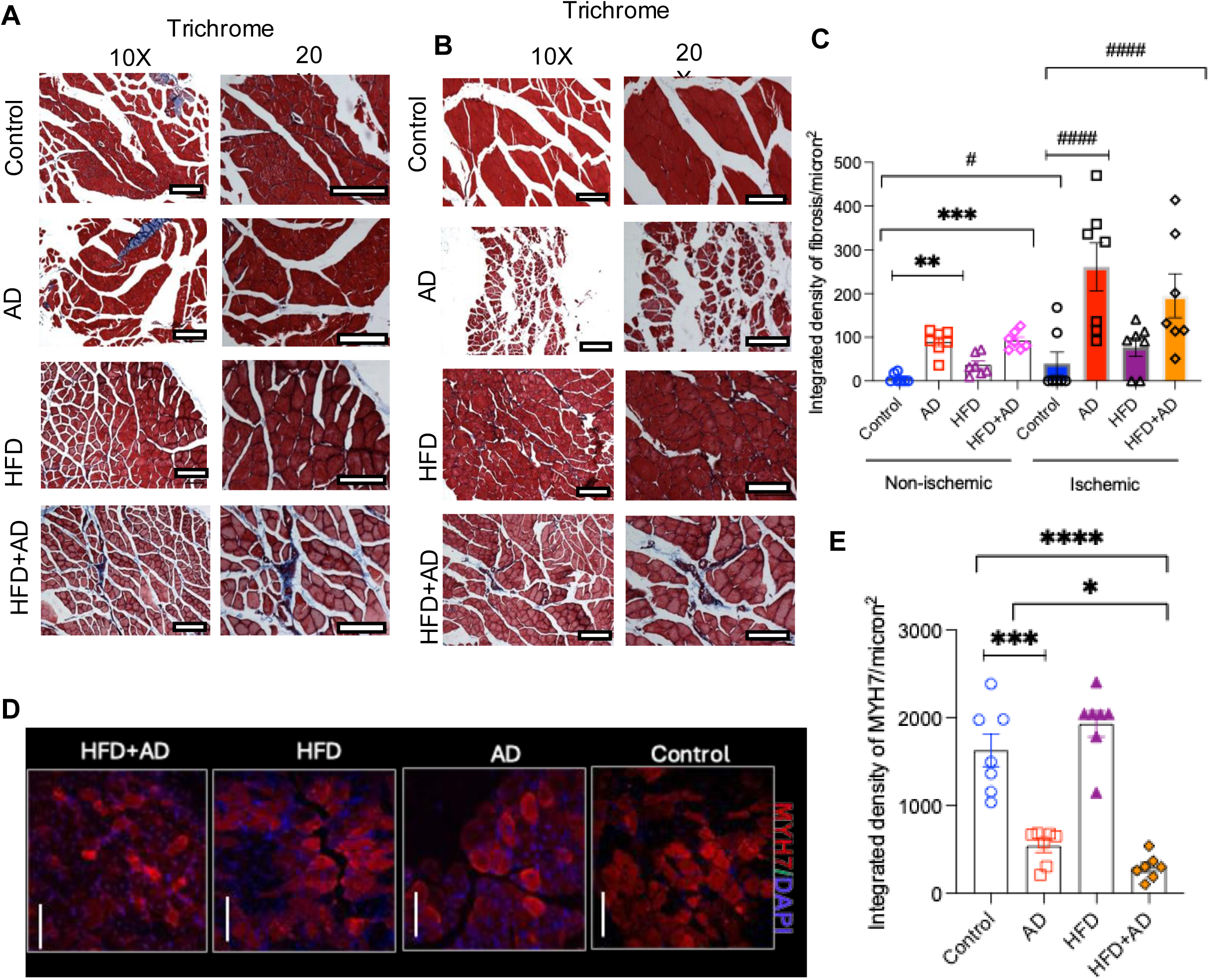
Muscle fibrosis in four groups of mice. **A)** Representative images from three random hpf images of trichrome-stained sections from non-ischemic muscle tissues are shown. (n = 7 mice/group), Scale bars = 100 microns. **B)** Representative images of trichrome-stained sections from ischemic soleus muscle tissues as above are shown. **C)** The muscle fibrosis was quantified in the trichrome-stained muscles, and three random trichrome-stained hpf images were used to quantify fibrosis. Fibrosis is represented as the integrated density of fibrosis normalized to the surface area. One-way ANOVA (P < 0.0001). In the non-ischemic group, compared to ND, the AD P = 0.016, and HFD+AD P = 0.004. In the ischemic group, one-way ANOVA (P < 0.0001). Compared to ND, the AD (P =0.0035), HFD (P =0.058), and HFD+AD (P =0.0179). Between the ischemic and non-ischemic ND groups P = 0.016. **D)** Representative images of ischemic soleus muscle stained from three random images (N= 7 mice/group) from the ligated limbs are shown. Scale bars = 100 microns. **E)** The MYH7 expression is represented as integrated density normalized to the surface area. One-way ANOVA P value of < 0.0001. Compared to the ND group, the AD group P = 0.0046) and the HFD+AD group P < 0.0001. Compared to the AD group, the HFD+AD group P = 0.028.

### Functional characterization of PAD in all groups of mice

Exercise is associated with skeletal muscle hyperemia, which is compromised in patients with PAD, resulting in vascular claudication^20^. This phenomenon is mimicked in mice as the reduction in post-compared to pre-exercise perfusion (flux ratio)^21^. The flux ratios were examined in all groups of mice at days 21 and 28 following HLI (**Figures 6A** and **6B**). In the ND group, there was an increase in flux ratio from 10% to 30% between days 21 and 28, consistent with an increase in perfusion and capillary density (**Figure 3A** and **3B**). At day 21, other groups showed a reduced flux ratio by 8 to 30%, with the lowest in mice on the HFD+AD diet. At day 28, while the flux ratio improved in all groups, yet compared to ND, the AD, HFD, and HFD+AD groups showed consistently lower flux ratio, with the lowest flux ratio was noted in the HFD+AD group (**Figure 6B**).

**Figure 6.**
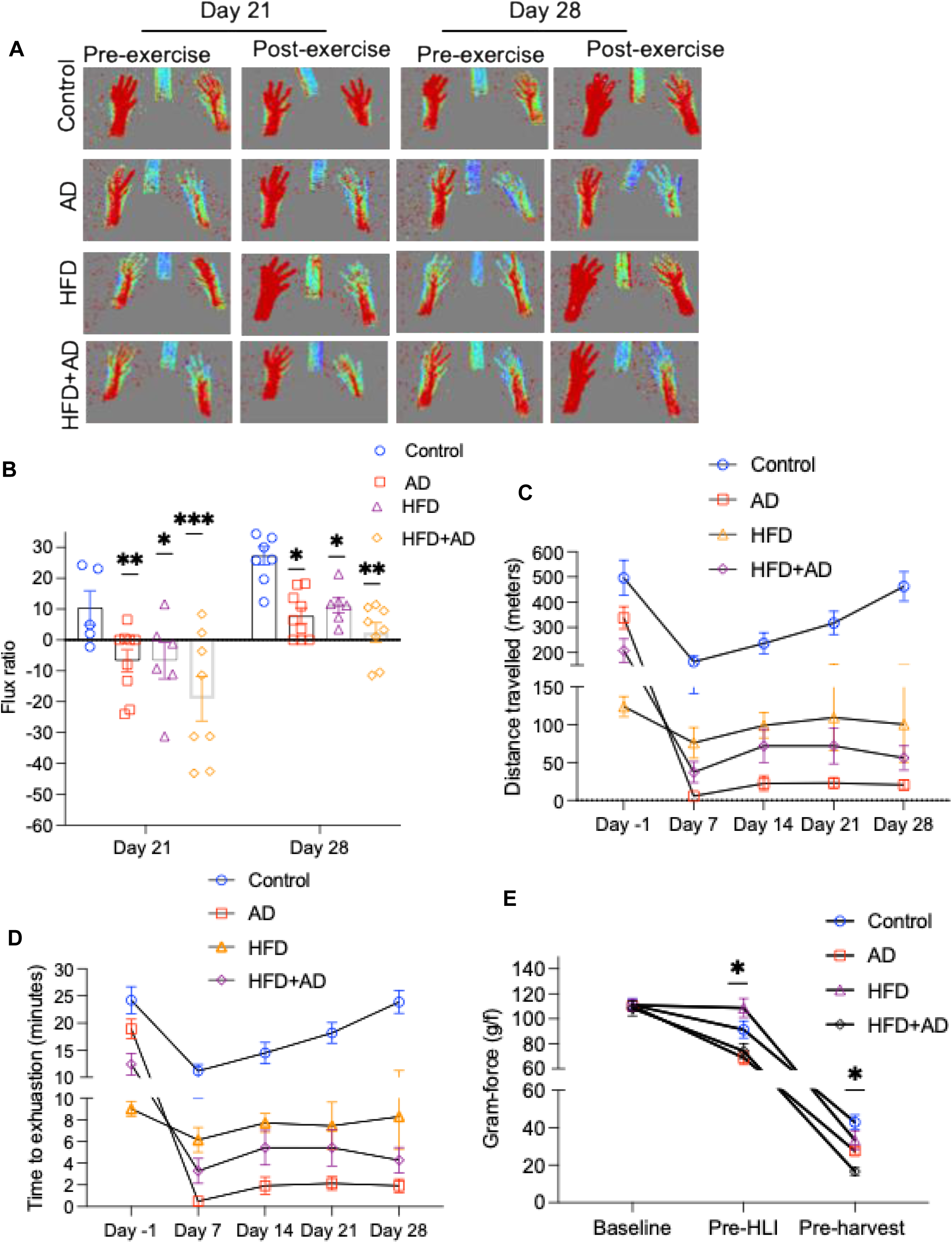
Functional characterization of the PAD model. **A)** Representative perfusion images of mice pre- and post-exercise at days 21 and 28 after HLI surgery are shown. **B)** The average flux ratios calculated as a difference between post- and pre-exercise perfusion are shown. Error bars = SEM. The mixed linear model analysis P < 0.0001. Tukey-Kramer adjustment for multiple comparisons: At day 21, Compared to the ND group, the AD, P = 0.038, HFD = 0.0449, and HFD+AD = 0.002. At day 28, compared to the ND group, AD P = 0.041, HFD = 0.038, and HFD+AD = 0.001. **C)** Distance traveled in meters at different time points are shown as average and error bars = SEM. A linear mixed model showed a significant interaction between time and groups P = 0.0001. The Tukey-Kramer adjustment was performed for multiple comparisons. At day 21, compared to the ND group, AD group P = 0.038, HFD P = 0.0449, and HFD +AD = 0.002. At 28 days, compared to ND, AD = 0.0001, HFD and HFD+AD < 0.0001. **D)** The average time to exhaustion among different groups of mice are shown. A mixed linear effect model demonstrated a significant interaction between time and group (P < 0.0001). At day 28, time to exhaustion was significantly reduced in all groups compared to ND, with P values as follows: AD = 0.0001, HFD+AD < 0.0001, and AD < 0.0001. **E)** The average grip strength of the mouse hind limb is depicted as grams/force at indicated time points. Error bars = SEM.

The distance traveled (meters) and time to exhaustion in all groups were recorded. Both these parameters showed reduction within seven days of HLI procedure, after which they improved in mice on ND (**Figures 6C** and **6D**). In the ND mice, the traveled distance increased from ∼180 meters in day seven to ∼ 220 meters on day 14, ∼ 300 meters in day 21 and ∼400 meters on day 28. However, for the mice on HFD and HFD+AD, the travel distances increased over time but were lower than the ND group of mice. The CKD mice showed consistently the lowest traveled distance. At the end of the experiment, all groups showed lower traveled distance compared to ND, and there was no significant difference between AD and HFD+AD groups.

Time to exhaustion followed a pattern similar to travel distance. The exhaustion time in ND mice was ∼ 12 minutes in day 7, ∼14 minutes in day 14, ∼ 18 minutes in day 21, and ∼ 25 minutes in day 28. In contrast, the exhaustion time remained consistently lower in other groups of mice. At the end of the experiment, there was no significant difference in the exhaustion time between AD and HFD+AD diet mice (**Figure 6D**). For the grip strength assessment, we measured the force generated in grams (g) at three-time points, as shown in **Figure 6E**. The baseline grip (pre-randomization) strength was similar in all groups. The mixed linear model showed a significant interaction between the time and different groups (pre-HLI P = 0.01, pre-harvest P = 0.03). Pre-HLI time point, compared to ND, the AD, and the HFD+AD groups, showed a 20-30% reduction (P = 0.023 and P = 0.031, respectively). At the end of the experiment, compared to the ND and AD groups, the HFD+AD group showed ∼ 15-30% reduction in the grip strength (P = 0.018 and P = 0.036, respectively), suggesting severe limb dysfunction in HFD+AD.

## Discussion

This work shows that a combination of AD and HFD diets in mice resulted in histopathological changes (IFTA and immune infiltration) and functional alterations in kidneys (low GFR). In addition, these mice displayed metabolic abnormalities characterized by hypercholesterolemia, glucose intolerance, hepatic steatosis and glomerulomegaly. The HFD+ AD mice also showed myocardial fibrosis and vascular abnormalities (reduced post-ischemic angiogenesis, compromised post-exercise hyperemia, lower grip strength, and endurance), all indicative of PAD. CKD mice showed earlier exhaustion and shorter traveled distance with a GFR of ∼40 µL/min at the end of 28 days, while HFD+AD mice showed similar levels of capillary rarefaction and limb dysfunction at a GFR ∼ double that of AD group. This point is clinically relevant as patients with CKM present with earlier CVD at a relatively preserved GFR^1–3, 5, 7^. Taken together, the HFD+AD model demonstrated several features of CKM syndrome. However, the HFD+ AD group did not show overt obesity, but presented evidence of glomerulomegaly, a renal phenotype associated with hyperfiltration due to type 2 diabetes and mild-moderate obesity in humans^22^. This phenotype may reflect elevated GFR observed during the earlier period of the HFD+ AD group.

CKM syndrome progresses through different stages, reflecting the progressive pathophysiology and increasing absolute CVD risk^1^. Stage 0 CKM includes individuals with no CVD, metabolic risk factors and normal renal function. Stage 1 CKM includes individuals with excess adipose tissue, dysfunctional adipose tissue (impaired glucose tolerance and hyperglycemia). Stage 2 includes individuals with metabolic risk factors (hypertriglyceridemia, hypertension, metabolic syndrome, or type 2 diabetes), moderate- to high-risk CKD, or both. Stage 3 includes individuals with subclinical CVD with overlapping CKM risk factors. Stage 4 includes individuals with clinical CVD (coronary heart disease, HF, stroke, peripheral artery disease, or atrial fibrillation) overlapping with CKM risk factors. The current animal model corrosponds to the CKM stage 3 or higher with overt CKD, and overt PAD. The earlier time points of this model can be explored to examine the previous stages of CKM and its progression through different stages.

This CKM model showed adipophilin expression in the liver. Adipophilin (ADRP/ADPH/PLIN2), a protein related to adipocyte differentiation, was first discovered in a mouse adipogenic cell line^17^. As a member of the perilipin (PAT protein family), it plays a crucial role in lipid metabolism and is an early marker of hepatic steatosis. Its downregulation in adenine mice is intriguing and warrants further investigation. The dietary modifications eased the generation of the model, and they have translational value, given that the diet has profound implications on the pathogenesis of CVD, metabolic abnormalities, and CKD.

The use of male mice and one type of CKD model represent the limitations of the current study. Our choice for the AD-induced CKD model was driven by its consistent induction of tubulointerstitial fibrosis, resulting in a rapid loss of GFR and accumulation of uremic toxins^10, 23, 24^. Further studies are needed to examine slowly progressive CKD (such as 5/6 nephrectomy), and metabolic models (genetic models) to induce CKM. While the choice of male mice in this model was driven by the prevalence of PAD in men^25^, women have a higher prevalence of PAD, especially from disadvantaged groups^8, 26^. Additional studies are needed to compare both sexes in mice examine the nature of ischemic myopathy.

CKM has profound public health implications and is a significant driver of disparities in CVD and outcomes. We hope this animal model will pave the pathway to deepen mechanistic understanding of cross-organ contribution to the pathogenesis of CKM and examine novel biomarkers and therapeutic targets.

## Acknowledgments

We thank Mike Kerber at the BUMC Imaging Core Facility for his assistance in image capture.

## Author Contributions

V.C.C. conceived and designed the project and edited the manuscript. S.L. assisted V.C.C. in concept design and writing manuscript; S.L, H.P, J.B and R.A. performed animal experiments, slide processing, staining, and image capture; M.B. performed renal histopathology analysis, H.C performed the statistical analysis, V.C.C and S.L. wrote the manuscript. All other authors edited the manuscript.

## Funding

This work was funded by the Center of Cross-Organ Vascular Pathology, DOM, BUSM, R01-HL166608, R01DK141016, R21-DK119740, R21-DK132784 (VCC), and AHA CAT-HD SFRN (VCC and SL), T32HL125232“Multidisciplinary Training Program in Cardiovascular Epidemiology (SL), Career Development Award, American Heart Association # 850917(W.Y.).

## Notes

### Competing Interest Statement

The authors have declared no competing interest.

